# ClassifyCNV: a tool for clinical annotation of copy-number variants

**DOI:** 10.1101/2020.07.20.213215

**Authors:** Tatiana A. Gurbich, Valery Vladimirovich Ilinsky

## Abstract

**Summary:** Copy-number variants (CNVs) are an important part of human genetic variation. They can be benign or can play a role in human disease by creating dosage imbalances and disrupting genes and regulatory elements. Accurate identification and clinical annotation of CNVs is essential when evaluating patients with neurodevelopmental disorders and congenital anomalies. Here, we present ClassifyCNV, a tool that implements the 2019 ACMG classification guidelines to assess CNV pathogenicity. ClassifyCNV uses genomic coordinates and CNV type as input and reports the clinical classification for each variant along with a classification score breakdown and a list of genes that could be important for variant interpretation. The tool is suitable for integration into NGS analysis pipelines and facilitates high-throughput CNV analysis.

**Availability and implementation:** ClassifyCNV is implemented in Python 3 and runs on UNIX, Linux and Mac OS X. The source code is available at https://github.com/Genotek/ClassifyCNV.

## Introduction

Copy-number variation is a form of structural genetic variation that involves a gain or loss of DNA segments. Copy-number variants (CNVs) are >50bp in size and can include a part of a gene, a whole gene, or a longer genomic region (Alkan et al., 2011). CNVs are associated with a number of genetic disorders, including autism spectrum disorders, neurodevelopmental disorders, and autoimmune diseases (Shaikh, 2017; Thapar & Cooper, 2013). With advancements in next-generation sequencing technology and an increasing availability of bioinformatics tools to analyze NGS data, clinical labs are now able to process and detect CNVs in batches of exomes, whole genomes, and gene panels. In order for the patients to receive an accurate diagnosis and appropriate care, it is essential to correctly determine the pathogenicity of variants.

In late 2019 ACMG released updated guidelines for clinical classification of CNVs (Riggs et al., 2020). Each CNV is classified into one of the following categories: benign, likely benign, a variant of uncertain significance, likely pathogenic, or pathogenic. The new guidelines take into account a wide range of CNV properties and allow for comprehensive analysis and accurate classification of variants. However, implementation of the guidelines on a large scale is challenging, as each CNV requires considerable time on the part of a clinician to obtain a final pathogenicity score. Although the new guidelines are intended for manual evaluation, computational analysis expedites the process and determines the impact of CNVs more efficiently. Available CNV annotation tools use criteria that are different from the new ACMG guidelines (Erikson et al., 2015; Ganel et al., 2017; Geoffroy et al., 2018), hence, a new computational approach is needed.

Here, we present ClassifyCNV, a command-line tool that allows for rapid high-throughput classification of CNVs in accordance with the latest ACMG guidelines.

## Implementation

ClassifyCNV is implemented in Python 3, runs on Linux, UNIX, and Mac OS X, and requires BEDTools v.2.27.1 or higher (Quinlan & Hall, 2010). It quickly processes large batches of CNVs and is easily integrated into other data analysis pipelines. Both the GRCh37 and the GRCh38 genome builds are supported.

ClassifyCNV accepts a BED file as input and requires the user to provide genomic coordinates and type (deletion or duplication) for each CNV. The tool then uses the criteria described in the ACMG scoring rubrics for copy number loss (http://cnvcalc.clinicalgenome.org/cnvcalc/cnv-loss) and gain (http://cnvcalc.clinicalgenome.org/cnvcalc/cnv-gain) to evaluate the clinical significance of the CNVs. Points are awarded for each evaluated section of the rubric. Clinical classification is calculated based on the total number of points assigned to a CNV (Supplementary methods, Tables S2.1 and S2.2).

To implement the assessment criteria ClassifyCNV analyzes the overlap between each CNV and protein-coding and noncoding genes, regulatory sequences, established and predicted dosage-sensitive genes and regions, and common genomic variants. A detailed description of the databases is provided in Supplementary methods, Table S1.

ClassifyCNV outputs a tab-delimited file that can be used by another pipeline in downstream analysis or evaluated by a clinician. For each variant ClassifyCNV reports the clinical classification, the total number of points, a breakdown of how the final pathogenicity score was determined, a list of established and predicted dosage-sensitive genes encompassed by the CNV, and a list of all protein-coding genes within the CNV. As some of the sections of the ACMG scoring rubrics require manual evaluation by a clinician, the information provided can be used to continue the evaluation if necessary.

## Performance testing

To test speed performance of ClassifyCNV, we used a set of copy number variants from the nstd102 study in ClinVar (38,488 unique CNVs in total) (Landrum et al., 2018). The run completed in less than 60 seconds on a 64-bit Linux virtual machine using two cores.

We used the same set of CNVs to evaluate the ClassifyCNV performance on clinical data. The pathogenic/ likely pathogenic variants and variants of uncertain significance had a high degree of concordance between the original ClinVar classification and the ClassifyCNV result (57% and 97.8% respectively). The majority of benign variants were classified as variants of uncertain significance (16,687 (87.7%)). 14,356 of these variants did not receive any points during the classification, indicating that the variants do contain genes or regulatory elements. However, the information about the genetic content within these variants was unavailable or did not strongly support reclassification of the variants from uncertain significance to benign or pathogenic. The results are shown in Supplementary results, Table S3.

To assess the concordance of the ClassifyCNV calls with the results of manual evaluation we obtained a list of 21 variants previously classified by the ACMG/ClinGen committee using the new guidelines (Supplementary results, Table S4). For all variants the automatic classification either matched the direction on the pathogenicity scale (i.e. assigned points towards pathogenicity for variants that were pathogenic according to the ACMG/ClinGen committee and assigned points towards the benign status for variants that were benign or likely benign) or left the number of points at 0, suggesting uncertain significance. For 71% of CNVs, the ClassifyCNV result matched the ACMG/ClinGen category. For variants automatically classified as uncertain, a manual evaluation of the published literature and patients’ family histories by a clinician was required to arrive at the final classification.

## Conclusions

ClassifyCNV is the first tool that automates the implementation of the updated ACMG guidelines to classify CNVs. It produces a rapid and reliable evaluation of variants and is suitable for high-throughput analysis. While a follow-up evaluation by a clinician may still be necessary for some variants, ClassifyCNV significantly expedites the evaluation process and allows for rapid filtration of benign variants. The tool is available at https://github.com/Genotek/ClassifyCNV.

## Acknowledgements

We thank Olesia Klimchuk for technical assistance and code review, Joshua Povich and Kirill Tsukanov for comments on the manuscript, and Alexandr Rakitko for testing the program and discussions that improved the project.

