## Supplementary data for "ClassifyCNV: a tool for clinical annotation of copy-number variants"

Supplementary Methods

1. Databases

The databases used to implement the 2019 ACMG criteria for clinical classification of copy-number variants (CNVs) (Riggs et al., 2020) are shown in Table S1. For each database we indicate which human genome build it is available for (hg19 or hg38). If a database is only available for one genome build, we used CrossMap v0.4.2 (Zhao et al., 2014) and the UCSC chain files, available from the UCSC genome browser (Kuhn et al., 2013), to lift over genomic coordinates between the genome builds.

All of the mentioned databases were converted to BED format and are available in the ClassifyCNV repository. We recommend that the local versions of the ClinGen databases are updated regularly by executing the update_clingen.sh script, which is available in the ClassifyCNV repository.

Table S1. Databases used by ClassifyCNV.

| Database | Genome build | Sections of the Copy Number Loss rubric where the database is used | Sections of the Copy Number Gain rubric where the database is used | Links and citations | Comment |
|---|---|---|---|---|---|
| RefGene | hg19, hg38 | 1, 3, 2E | 1, 3, 2D, 2E, 2F, 2G | http://hgdownload.cse.ucsc.edu/goldenPath/hg19/database/refGene.txt.gz<br><br>http://hgdownload.cse.ucsc.edu/goldenPath/hg38/database/refGene.txt.gz<br><br>(O’Leary et al., 2016) | Only one transcript per gene was kept. A MANE Select transcript was used if available. Otherwise, the longest transcript was used. |
| Promoters | hg19, hg38 | 1 | 1 | - | The promoter database was generated by obtaining the coordinates of the 500bp region directly upstream of every gene in the refGene |

| | | | | | |
|---|---|---|---|---|---|
| | | | | | database. |
| VISTA enhancers | hg19 | 1 | 1 | https://enhancer.lbl.gov/cgi-bin/imagedb3.pl?page_size=10((0;show=1;search.result=yes;page=1;form=search;search.form=no;action=search;search.sequence=1<br><br>(Visel et al., 2007) | The database was pre-parsed to only keep human enhancers.<br><br>This database is combined with FANTOM5 enhancers and enhancers from the Ensembl regulatory build to make the Enhancers.3sources.merged.bed file. |
| FANTOM5 enhancers | hg19, hg38 | 1 | 1 | https://fantom.gsc.riken.jp/5/datafiles/latest/extra/Enhancers/human_permissive_enhancers_phase_1_and_2.bed.gz<br><br>https://fantom.gsc.riken.jp/5/datafiles/reprocessed/hg38_latest/extra/enhancer/F5.hg38.enhancers.bed.gz<br><br>(Andersson et al., 2014) | This database is combined with VISTA enhancers and enhancers from the Ensembl regulatory build to make the Enhancers.3sources.merged.bed file. |
| Ensembl regulatory build | hg38 | 1 | 1 | ftp://ftp.ensembl.org/pub/release-100/regulation/homo_sapiens/homo_sapiens.GRCh38.Regulatory_Build.regulatory_features.20190329.gff.gz<br><br>(Zerbino et al., 2015) | Only enhancers were extracted from the database.<br><br>This database is combined with VISTA enhancers and FANTOM5 enhancers to make the Enhancers.3sources.merged.bed file. |
| refGene gene features (5’UTR, 3’UTR, exons, cds) | hg19, hg38 | 2C, 2D, 2E | 2E | http://genome.ucsc.edu/cgi-bin/hgTables<br><br>(Kuhn et al., 2013) | Gene feature coordinates were obtained for protein-coding genes. Only one transcript per gene was used (MANE Select if available, the longest one otherwise). |
| ClinGen haploinsufficient and triplosensitive genes and curated regions | hg19, hg38 | 2A-2G | 2A-2H | ftp://ftp.clinicalgenome.org/<br><br>(Rehm et al., 2015) | The script that downloads and parses the databases (update_clingen.sh) is included in the repository. |
| DECIPHER HI Predictions Version3 | hg38 | 2H | | https://decipher.sanger.ac.uk/files/downloads/HI_Predictions_Version3.bed.gz<br><br>(Firth et al., 2009) | |
| ExAC pLI scores | hg19 | 2H | | ftp://ftp.broadinstitute.org/pub/ExAC_release/release1/manuscript_data/forweb_cleaned_exac_r03_march16_z_data_pLI.txt.gz | |

| | | | | | |
|---|---|---|---|---|---|
| | | | | (Lek et al., 2016) | |
| the pLoF observed/expected upper fraction (LOEUF) | hg19 | 2H | | gs://gnomad-public/release/2.1.1/constraint/gnomad.v2.1.1.lof_metrics.by_gene.txt.bgz<br><br>(Collins et al., 2020) | |
| DGV Gold Standard Variants (2016-05-15) | hg19, hg38 | 4O | 4O | http://dgv.tcag.ca/dgv/docs/DGV.GS.March2016.50percent.GainLossSep.Final.hg19.gff3<br><br>http://dgv.tcag.ca/dgv/docs/DGV.GS.hg38.gff3<br><br>(MacDonald et al., 2014) | Included in populations_freqs.bed |
| gnomAD structural variant frequencies | hg19 | 4O | 4O | https://storage.googleapis.com/gnomad-public/papers/2019-sv/gnomad_v2.1_sv.sites.bed.gz<br><br>(Collins et al., 2020) | Included in populations_freqs.bed |

2. Rubric implementation

To assess the genomic content of each variant, ClassifyCNV checks for a full or partial (≥ 1bp) overlap with protein-coding and noncoding genes, as well as enhancers and promoters. It also tracks the number of protein-coding genes that are fully or partially overlapped by each CNV. To assess whether any established dosage-sensitive genes or regions are included and what effect the deletion or duplication might have on their expression, each CNV is evaluated against a set of curated haploinsufficient and triplosensitive genes and genomic regions obtained from ClinGen (Rehm et al., 2015). A score of ‘3’ is required for a gene or genomic region to be considered haploinsufficient or triplosensitive. For partially overlapped dosage-sensitive genes ClassifyCNV evaluates which regions within the gene are involved as per ACMG guidelines. If a deletion does not encompass genes or regions that are known to be haploinsufficient, ClassifyCNV checks whether haploinsufficiency is predicted for any genes within the deletion.

To satisfy this condition, a gene is required to have a DECIPHER HI index ≤ 10% (Firth et al., 2009), a gnomAD pLI score ≥ 0.9 and the upper bound of the observed/expected confidence interval <0.35 (Karczewski et al., 2020). Finally, to assess whether the CNV is likely to be benign, ClassifyCNV obtains the population frequencies of similar variants from DGV (MacDonald et al., 2014) and gnomAD (Collins et al., 2020). For each analyzed CNV that does not contain known dosage-sensitive genes or genomic regions, the population frequencies of known overlapping CNVs are extracted. An overlap of at least 80% of the query CNV length is required. If multiple known variants overlap the CNV, their average population frequency is calculated. A CNV is considered common if its population frequency is > 1%.

ClassifyCNV continues through the end of the rubric for all CNVs, including the ones where a benign or pathogenic classification is determined before all of the conditions in the rubric have been evaluated.

The criteria that are implemented in ClassifyCNV are listed in Table S2.1 for copy number losses and in Table S2.2 for copy number gains.

Table S2.1. Implementation of the Copy Number Loss rubric

**Section 1: Initial assessment of genomic content**

| Evidence type | Evidence | The number of points suggested by the ACMG/Clingen guidelines | The number of points ClassifyCNV assigns if the condition is satisfied | Implementation |
|---|---|---|---|---|
| Copy-number loss content | **1A.** Contains protein-coding or other known functionally important elements. | 0 | 0 | Implemented |
| | **1B.** Does NOT contain protein-coding or any known | −0.60 | −0.60 | Implemented |

| | |
|---|---|
| | functionally important elements. |

**Section 2: Overlap with established/predicted haploinsufficiency (HI) or established benign genes/genomic regions** ***(Skip to section 3 if your copy-number loss DOES NOT overlap these types of genes/regions)***

| | | | | |
|---|---|---|---|---|
| Overlap with ESTABLISHED HI genes or genomic regions and consideration of reason for referral | **2A.** Complete overlap of an established HI gene/genomic region. | 1 | 1 | Implemented |
| | **2B.** Partial overlap of an established HI genomic region<br>• The observed CNV does NOT contain the known causative gene or critical region for this established HI genomic region OR<br>• Unclear if known causative gene or critical region is affected OR<br>• No specific causative gene or critical region has been established for this HI genomic region | 0 | 0 | Implemented |
| | **2C.** Partial overlap with the 5' end of an established HI gene (3' end of the gene not involved)… | | See categories below | |
| | **2C-1.** …and coding sequence is involved | 0.90 (Range: 0.45 to 1.00) | 0.90 | Implemented |
| | **2C-2.** …and only the 5' UTR is involved | 0 (Range: 0 to 0.45) | 0 | Implemented |
| | **2D.** Partial overlap with the 3' end of an established HI gene (5' end of the gene not involved)… | | See categories below | |
| | **2D-1.** …and only the 3' untranslated region is involved. | 0 | 0 | Implemented |
| | **2D-2.** …and only the last exon is involved. Other established pathogenic variants have been reported | 0.90 (Range: 0.45 to 0.90) | 0.30 | Partially implemented.<br><br>0.30 points are assigned in all cases |

| | | | | |
|---|---|---|---|---|
| | in this exon. | | | where only the last exon is involved regardless of whether there are established pathogenic variants in this exon. |
| | **2D-3.** …and only the last exon is involved. No other established pathogenic variants have been reported in this exon. | 0.30 (Range: 0 to 0.45) | 0.30 | |
| | **2D-4**. …and it includes other exons in addition to the last exon. Nonsense-mediated decay is expected to occur. | 0.90 (Range: 0.45 to 1.00) | 0.90 | Implemented |
| | **2E.** Both breakpoints are within the same gene (intragenic CNV; gene-level sequence variant). | See ClinGen SVI working group PVS1 specifications<br>• PVS1 = 0.90 *(Range: 0.45 to 0.90)*<br>• PVS1_Strong = 0.45 *(Range: 0.30 to 0.90)*<br>• PVS1_Moderate *or* PM4 (in-frame indels) = 0.30 *(Range: 0.15 to 0.45)*<br>• PVS1_Supporting = 0.15 *(Range: 0 to 0.30)*<br>• N/A = No points, but continue evaluation | • PVS1 = 0.90<br>• N/A = No points, but continue evaluation | Partially implemented.<br><br>If the --precise flag is on, points are assigned for intragenic variants in biologically-relevant transcripts that disrupt the reading frame and are predicted to lead to nonsense-mediated decay. No points are assigned for other types of intragenic deletions. |
| | **2F.** Completely contained within an established benign CNV region. | −1 | −1 | Implemented |
| | **2G**. Overlaps an established benign CNV, but includes additional genomic material. | 0 | 0 | Implemented |
| | **2H.** Two or more HI predictors suggest that AT LEAST ONE gene in the interval is HI. | 0.15 | 0.15 | Implemented |

**Section 3: Evaluation of gene number**

| | | | | |
|---|---|---|---|---|
| Number of protein-coding RefSeq genes wholly or | **3A.** 0–24 genes | 0 | 0 | Implemented (Note: if genes belong to the same gene |

| | | | | |
|---|---|---|---|---|
| partially included in the copy-number loss | | | | family, each gene within the family is counted) |
| | **3B.** 25–34 genes | 0.45 | 0.45 | Implemented (Note: if genes belong to the same gene family, each gene within the family is counted) |
| | **3C.** 35+ genes | 0.90 | 0.90 | Implemented (Note: if genes belong to the same gene family, each gene within the family is counted) |

**Section 4: Detailed evaluation of genomic content using cases from published literature, public databases, and/or internal lab data*(Skip to section 5 if either your CNV overlapped with an established HI gene/region in section 2, OR there have been no reports associating either the CNV or any genes within the CNV with human phenotypes caused by loss of function [LOF] or copy-number loss)***

| | | | | |
|---|---|---|---|---|
| Individual case evidence — de novo occurrences | Reported proband (from literature, public databases, or internal lab data) has either:<br>• A complete deletion of or a LOF variant within gene encompassed by the observed copy-number loss OR<br>• An overlapping copy-number loss similar in genomic content to the observed copy-number loss AND… | See categories below | | |
| | **4A**. …the reported phenotype is highly specific and relatively unique to the gene or genomic region, | Confirmed de novo: 0.45 points each Assumed de novo: 0.30 points each (range: 0.15 to 0.45). 0.90 (total) | – | Not implemented |
| | **4B.** …the reported phenotype is consistent with the gene/genomic region, is highly specific, but not necessarily unique to the gene/genomic region. | Confirmed de novo: 0.30 points each Assumed de novo: 0.15 point each (range: 0 to 0.45) | – | Not implemented |
| | **4C.** …the reported phenotype is consistent with the | Confirmed de novo: 0.15 point each Assumed de novo: | – | Not implemented |

| | | | | |
|---|---|---|---|---|
| | gene/genomic region, but not highly specific and/or with high genetic heterogeneity. | 0.10 point each (range: 0 to 0.30) | | |
| Individual case evidence — inconsistent phenotype | **4D.** …the reported phenotype is NOT consistent with what is expected for the gene/genomic region or not consistent in general. | 0 points each (range: 0 to −0.30). −0.30 (total) | – | Not implemented |
| Individual case evidence — unknown inheritance | **4E.** Reported proband has a highly specific phenotype consistent with the gene/genomic region, but the inheritance of the variant is unknown. | 0.10 points each (range: 0 to 0.15). 0.30 (total) | – | Not implemented |
| Individual case evidence — segregation among similarly affected family members | **4F.** 3–4 observed segregations | 0.15 (0.45 max) | – | Not implemented |
| | **4G.** 5–6 observed segregations | 0.30 | – | Not implemented |
| | **4H.** 7 or more observed segregations | 0.45 | – | Not implemented |
| Individual case evidence — nonsegregations | **4I.** Variant is NOT found in another individual in the proband's family AFFECTED with a consistent, specific, well-defined phenotype (no known phenocopies). | −0.45 points per family (range: 0 to −0.45). −0.90 (total) | – | Not implemented |
| | **4J.** Variant IS found in another individual in the proband's family UNAFFECTED with the specific, well-defined phenotype observed in the proband. | −0.30 points per family (range: 0 to −0.30). −0.90 (total) | – | Not implemented |
| | **4K.** Variant IS found in another individual | −0.15 points per family (range: 0 to | – | Not implemented |

| | | | | |
|---|---|---|---|---|
| | in the proband's family UNAFFECTED with the nonspecific phenotype observed in the proband. | −0.15). −0.30 (total) | | |
| Case–control and population evidence | **4L.** Statistically significant increase amongst observations in cases (with a consistent, specific, well-defined phenotype) compared with controls. | 0.45 per study (range: 0 to 0.45 per study). 0.45 (total) | – | Not implemented |
| | **4M.** Statistically significant increase amongst observations in cases (without a consistent, nonspecific phenotype OR unknown phenotype) compared with controls. | 0.30 per study (range: 0 to 0.30 per study). 0.45 (total) | – | Not implemented |
| | **4N.** No statistically significant difference between observations in cases and controls. | −0.90 (per study) (range: 0 to −0.90 per study). −0.90 (total) | – | Not implemented |
| | **4O.** Overlap with common population variation. | −1 (range: 0 to −1) | −1 | Implemented |
| **Section 5: Evaluation of inheritance pattern/family history for patient being studied** | | | | |
| Observed copy-number loss is de novo | **5A.** *Use appropriate category from de novo scoring section in section 4.* | *Use de novo scoring categories from section 4 (4A–4D) to determine score (0.45 max)* | – | Not implemented |
| Observed copy-number loss is inherited | **5B.** Patient with specific, well-defined phenotype and no family history. CNV is inherited from an apparently unaffected parent. | −0.30 (range: 0 to −0.45) | – | Not implemented |
| | **5C.** Patient with nonspecific phenotype and no family history. CNV is inherited from an | −0.15 (range: 0 to −0.30) | – | Not implemented |

| | | | | |
|---|---|---|---|---|
| | apparently unaffected parent. | | | |
| | **5D.** CNV segregates with a consistent phenotype observed in the patient's family. | *Use segregation scoring categories from section 4 (4F–4H) to determine score (0.45 max)* | – | Not implemented |
| Observed copy-number loss — nonsegregations | **5E.** *Use appropriate category from nonsegregation section in section 4*. | *Use nonsegregation scoring categories from section 4 (4I–4K) to determine score (–0.45 max)* | – | Not implemented |
| Other | **5F.** Inheritance information is unavailable or uninformative. | 0 | – | Not implemented |
| | **5G.** Inheritance information is unavailable or uninformative. The patient phenotype is nonspecific, but is consistent with what has been described in similar cases. | 0.10 (range: 0 to 0.15) | – | Not implemented |
| | **5H.** Inheritance information is unavailable or uninformative. The patient phenotype is highly specific and consistent with what has been described in similar cases. | 0.30 (range: 0 to 0.30) | – | Not implemented |

Table S2.2. Implementation of the Copy Number Gain rubric.

**Section 1: Initial assessment of genomic content**

| **Evidence type** | **Evidence** | **The number of points suggested by the ACMG/Clingen guidelines** | **The number of points ClassifyCNV assigns if the condition is satisfied** | **Implementation** |
|---|---|---|---|---|
| Copy-number gain content | **1A.** Contains protein-coding or other known functionally important elements. | 0 | 0 | Implemented |

| | | | | |
|---|---|---|---|---|
| | **1B.** Does NOT contain protein-coding or any known functionally important elements. | −0.60 | −0.60 | Implemented |

**Section 2: Overlap with established triplosensitive (TS), haploinsufficient (HI), or benign genes or genomic regions** ***(Skip to section 3 if the copy-number gain DOES NOT overlap these types of genes/regions)***

| | | | | |
|---|---|---|---|---|
| Overlap with ESTABLISHED TS genes or genomic regions | **2A.** Complete overlap; the TS gene or minimal critical region is fully contained within the observed copy-number gain. | 1 | 1 | Implemented |
| | **2B.** Partial overlap of an established TS region<br>• The observed CNV does NOT contain the known causative gene or critical region for this established TS genomic region OR<br>• Unclear if known causative gene or critical region is affected OR<br>• No specific causative gene or critical region has been established for this TS genomic region | 0 | 0 | Implemented |
| Overlap with ESTABLISHED benign copy-number gain genes or genomic regions | **2C.** Identical in gene content to the established benign copy-number gain. | −1 | −1 | Implemented |
| | **2D.** Smaller than established benign copy-number gain, breakpoint(s) does not interrupt protein-coding genes. | −1 | −1 | Implemented (only when the --precise flag is on) |
| | **2E.** Smaller than established benign copy-number gain, breakpoint(s) potentially interrupts protein-coding gene. | 0 | 0 | Implemented |
| | **2F.** Larger than known benign copy-number gain, does | −1 (range: 0 to −1.00) | −1 | Implemented |

| | | | | |
|---|---|---|---|---|
| | not include additional protein-coding genes. | | | |
| | **2G.** Overlaps a benign copy-number gain but includes additional genomic material. | 0 | 0 | Implemented |
| Overlap with ESTABLISHED HI gene(s) | **2H.** HI gene fully contained within observed copy-number gain. | 0 | 0 | Implemented |
| Breakpoint(s) within ESTABLISHED HI genes | **2I.** Both breakpoints are within the same gene (gene-level sequence variant, possibly resulting in loss of function [LOF]). | See ClinGen SVI working group PVS1 specifications<br>• PVS1 = 0.90 (Range: 0.45 to 0.90)<br>• PVS1_Strong = 0.45 (Range: 0.30 to 0.90)<br>• N/A = 0 (Continue evaluation) | – | Not implemented |
| | **2J.** One breakpoint is within an established HI gene, patient's phenotype is either inconsistent with what is expected for LOF of that gene OR unknown. | 0 | 0 | Implemented |
| | **2K.** One breakpoint is within an established HI gene, patient's phenotype is highly specific and consistent with what is expected for LOF of that gene. | 0.45 | – | Not implemented |
| Breakpoints within other gene(s) | **2L.** One or both breakpoints are within gene(s) of no established clinical significance. | 0 | 0 | Implemented |

**Section 3: Evaluation of gene number**

| | | | | |
|---|---|---|---|---|
| Number of protein-coding RefSeq genes wholly or partially included in the copy-number gain | **3A.** 0–34 genes | 0 | 0 | Implemented (Note: if genes belong to the same gene family, each gene within the family is counted) |

| | | | | |
|---|---|---|---|---|
| | **3B.** 35–49 genes | 0.45 | 0.45 | Implemented (Note: if genes belong to the same gene family, each gene within the family is counted) |
| | **3C.** 50+ genes | 0.90 | 0.90 | Implemented (Note: if genes belong to the same gene family, each gene within the family is counted) |

**Section 4: Detailed evaluation of genomic content using cases from published literature, public databases, and/or internal lab data*(Note: If there have been no reports associating either the copy-number gain or any of the genes therein with human phenotypes caused by triplosensitivity, skip to section 5)***

| | | | | |
|---|---|---|---|---|
| Individual case evidence — de novo occurrences | Reported proband (from literature, public databases, or internal lab data) has either:<br>• complete duplication of one or more genes within the observed copy-number gain OR<br>• an overlapping copy-number gain similar in genomic content to the observed copy-number gain AND… | See categories below | | |
| | **4A.** …the reported phenotype is highly specific and relatively unique to the gene or genomic region. | Confirmed de novo: 0.45 points each Assumed de novo: 0.30 points each (range: 0.15 to 0.45). 0.90 (total) | – | Not implemented |
| | **4B.** …the reported phenotype is consistent with the gene/genomic region, is highly specific, but not necessarily unique to the gene/genomic region. | Confirmed de novo: 0.30 points each Assumed de novo: 0.15 point each (range: 0 to 0.45) | – | Not implemented |
| | **4C.** …the reported phenotype is consistent with the gene/genomic region, but not highly specific and/or with high genetic heterogeneity. | Confirmed de novo: 0.15 point each Assumed de novo: 0.10 point each (range: 0 to 0.30) | – | Not implemented |

| | | | | |
|---|---|---|---|---|
| Individual case evidence — inconsistent phenotype | **4D.** …the reported phenotype is NOT consistent with the gene/genomic region or not consistent in general. | 0 points each (range: 0 to −0.30). −0.30 (total) | – | Not implemented |
| Individual case evidence — unknown inheritance | **4E.** Reported proband has a highly specific phenotype consistent with the gene/genomic region, but the inheritance of the variant is unknown. | 0.10 points each (range: 0 to 0.15). 0.30 (total) | – | Not implemented |
| Individual case evidence — segregation among similarly affected family members | **4F.** 3–4 observed segregations | 0.15 (0.45 max) | – | Not implemented |
| | **4G.** 5–6 observed segregations | 0.30 | – | Not implemented |
| | **4H.** 7 or more observed segregations | 0.45 | – | Not implemented |
| Individual case evidence — nonsegregations | **4I.** Variant is NOT found in another individual in the proband's family AFFECTED with a consistent, specific, well-defined phenotype (no known phenocopies). | −0.45 points per family (range: 0 to −0.45). −0.90 (total) | – | Not implemented |
| | **4J.** Variant IS found in another individual in the proband's family UNAFFECTED with the specific, well-defined phenotype observed in the proband. | −0.30 points per family (range: 0 to −0.30). −0.90 (total) | – | Not implemented |
| | **4K.** Variant IS found in another individual in the proband's family UNAFFECTED with the nonspecific phenotype observed in the proband. | −0.15 points per family (range: 0 to −0.15). −0.30 (total) | – | Not implemented |

| | | | | |
|---|---|---|---|---|
| Case–control and population evidence | **4L.** Statistically significant increase amongst observations in cases (with a consistent, specific, well-defined phenotype) compared with controls. | 0.45 per study (range: 0 to 0.45 per study). 0.45 (total) | – | Not implemented |
| | **4M.** Statistically significant increase amongst observations in cases (without a consistent, nonspecific phenotype OR unknown phenotype) compared with controls. | 0.30 per study (range: 0 to 0.30 per study). 0.45 (total) | – | Not implemented |
| | **4N.** No statistically significant difference between observations in cases and controls. | −0.90 (per study) (range: 0 to −0.90 per study). −0.90 (total) | – | Not implemented |
| | **4O.** Overlap with common population variation. | −1 (range: 0 to −1) | −1 | Implemented |
| **Section 5: Evaluation of inheritance pattern/family history for patient being studied** | | | | |
| Observed copy-number gain is de novo | **5A.** *Use appropriate category from de novo scoring section in section 4.* | *Use de novo scoring categories from section 4 (4A–4D) to determine score (0.45 max)* | – | Not implemented |
| Observed copy-number loss is inherited | **5B.** Patient with specific, well-defined phenotype and no family history. Copy-number gain is inherited from an apparently unaffected parent. | −0.30 (range: 0 to −0.45) | – | Not implemented |
| | **5C.** Patient with nonspecific phenotype and no family history. Copy-number gain is inherited from an apparently unaffected parent. | −0.15 (range: 0 to −0.30) | – | Not implemented |
| | **5D.** CNV segregates with consistent | *Use segregation scoring categories* | – | Not implemented |

| | | | | |
|---|---|---|---|---|
| | phenotype observed in the patient's family. | *from section 4 (4F–4H) to determine score (0.45 max)* | | |
| Observed copy-number gain — nonsegregations | **5E.** *Use appropriate category from nonsegregation section in section 4.* | *Use nonsegregation scoring categories from section 4 (4I–4K) to determine score (–0.45 max)* | – | Not implemented |
| | **5F.** Inheritance information is unavailable or uninformative. | 0 | – | Not implemented |
| | **5G.** Inheritance information is unavailable or uninformative. The patient phenotype is nonspecific, but is consistent with what has been described in similar cases. | 0.10 (range: 0 to 0.15) | – | Not implemented |
| | **5H.** Inheritance information is unavailable or uninformative. The patient phenotype is highly specific and consistent with what has been described in similar cases. | 0.15 (range: 0 to 0.30) | – | Not implemented |

Supplementary Results

To test the ClassifyCNV performance on clinical data, we obtained a set of 17,683 duplications and 20,805 deletions from the nstd102 study in ClinVar (Landrum et al., 2018). We used the hg19 coordinates and ran ClassifyCNV using the --precise flag, thus treating the CNV coordinates as exact. For CNVs for which precise coordinates were unknown, we used the inner coordinates. The ClinVar variants were obtained from studies published prior to 2019 and, therefore, classified before the current ACMG guidelines were released. The comparison of ClinVar and ClassifyCNV classifications is shown in Table S3.

The pathogenic/ likely pathogenic variants and variants of uncertain significance had a high degree of concordance between the original ClinVar classification and the ClassifyCNV result (57% and 97.8%, respectively). The majority of benign variants were classified as variants of uncertain significance (16,687 (87.7%)). 14,356 of these variants did not receive any points during the classification, indicating that the variants do contain genes or regulatory elements. However, the information about the genetic content within these variants was unavailable or did not strongly support reclassification of the variants from uncertain significance to benign or pathogenic. Although points were not awarded and a follow-up evaluation by a clinician may be necessary, ClassifyCNV completed the evaluation of gene content, dosage-sensitivity, and population frequencies of these variants and outputted a list of genes of interest, thus facilitating follow-up evaluations.

ClassifyCNV errs on the side of caution when moving a variant between categories, as advised by the new ACMG guidelines. Therefore, if convincing data are not available, a CNV is likely to remain a variant of uncertain significance. Additionally, since the classification parameters used by ClassifyCNV are different from the parameters used prior to the release of the 2019 ACMG guidelines, we do not expect full concordance even when evaluating variants manually.

Table S3. ClassifyCNV performance on ClinVar data.

| **ClinVar Classification** | | **ClassifyCNV Classification** | |
|---|---|---|---|
| *Classification* | *Count* | *Classification* | *Count (percentage)* |
| Pathogenic/ Likely pathogenic | 6,780 | Pathogenic/ Likely pathogenic | 3,865 (57%) |
| | | Uncertain significance | 2,902 (42.8%) |
| | | Benign/ Likely benign | 13 (0.2%) |

| | | | |
|---|---|---|---|
| Benign/ Likely benign | 19,026 | Benign/ Likely benign | 2,246 (11.8%) |
| | | Uncertain significance | 16,687 (87.7%) |
| | | Pathogenic/ Likely pathogenic | 93 (0.5%) |
| Uncertain significance | 12,682 | Uncertain significance | 12,394 (97.8%) |
| | | Benign/ Likely benign | 69 (0.5%) |
| | | Pathogenic/ Likely pathogenic | 219 (1.7%) |

To test the concordance between the ClassifyCNV calls and the manual evaluation by ACMG/Clingen we obtained four manually evaluated CNVs from Riggs et al 2020 and 17 examples of CNV evaluation from the ClinGen website (https://clinicalgenome.org/tools/cnv-webinar/examples/). ClassifyCNV was executed with the --precise flag on (CNV breakpoints were presumed to be accurate). The results are shown in Table S4. In all cases where ClassifyCNV determined that the pathogenicity of a variant was uncertain while the manual evaluation arrived at a benign/likely benign or a pathogenic classification, the difference was due to the results of manual evaluation of published literature, population data and patient's family history.

Table S4. Comparison of ClassifyCNV calls to the results of manual annotation by ACMG/Clingen. All coordinates are hg19. For 71% of CNVs, the ClassifyCNV result matched the ACMG/ClinGen result category exactly. For the remaining cases, a manual follow-up evaluation of published literature, population data and patients' family histories was required to arrive at the final classification. In no cases did the ClassifyCNV result differ from the ACMG/ClinGen result in a manner that would lead to an erroneous clinical interpretation.

| Chromosome | Start position | End position | CNV type | ACMG classification | ClassifyCNV classification | ClassifyCNV score |
|---|---|---|---|---|---|---|
| chr19 | 43,242,796 | 43,741,310 | DEL | Benign | Benign | <−1 |
| chr15 | 30,507,853 | 30,807,921 | DEL | Benign | Likely benign | −0.9 |
| chr13 | 20,812,000 | 21,012,000 | DEL | Benign | Uncertain | 0 |
| chr2 | 45,408,934 | 45,976,420 | DUP | Benign | Uncertain | 0 |
| chr4 | 69,373,811 | 69,491,113 | DEL | Benign | Likely benign | −0.9 |
| chr19 | 18,291,753 | 18,311,626 | DUP | Benign | Likely benign | −0.9 |
| chr1 | 1,379,519 | 1,435,227 | DEL | Likely benign | Uncertain | 0 |
| chr22 | 18,912,231 | 21,465,672 | DUP | Pathogenic | Pathogenic | >1 |
| chr12 | 12,864,101 | 14,983,330 | DEL | Pathogenic | Pathogenic | 1 |
| chr11 | 45,904,399 | 46,480,747 | DEL | Pathogenic | Uncertain | 0.15 |
| chr18 | 53,049,652 | 53,134,356 | DEL | Pathogenic | Likely pathogenic | 0.9 |
| chr17 | 553,250 | 1,353,250 | DEL | Pathogenic | Uncertain | 0.15 |
| chr22 | 31,645,547 | 32,807,482 | DEL | Pathogenic | Uncertain | 0 |
| chr9 | 108,597,937 | 111,269,478 | DEL | Pathogenic | Pathogenic | >1 |
| chrX | 23,223,505 | 23,660,309 | DEL | Pathogenic | Pathogenic | 1 |
| chr5 | 125,989,631 | 126,295,396 | DUP | Pathogenic | Pathogenic | 1 |
| chr12 | 27,715,516 | 29,628,080 | DEL | Uncertain | Uncertain | 0 |
| chr17 | 41,784,108 | 42,438,203 | DUP | Uncertain | Uncertain | 0 |
| chr22 | 40,654,201 | 40,659,533 | DEL | Uncertain | Uncertain | 0 |
| chr3 | 190,380,498 | 191,783,134 | DEL | Uncertain | Uncertain | 0 |
| chr17 | 13,029,888 | 14,707,559 | DUP | Uncertain | Uncertain | 0 |
